# Viral Quasispecies Reconstruction via Correlation Clustering

**DOI:** 10.1101/096768

**Authors:** Somsubhra Barik, Shreepriya Das, Haris Vikalo

## Abstract

RNA viruses are characterized by high mutation rates that give rise to populations of closely related viral genomes, the so-called viral quasispecies. The underlying genetic heterogeneity occurring as a result of natural mutation-selection process enables the virus to adapt and proliferate in face of changing conditions over the course of an infection. Determining genetic diversity (i.e., inferring viral haplotypes and their proportions in the population) of an RNA virus is essential for the understanding of its origin and mutation patterns, and the development of effective drug treatments. In this paper we present QSdpR, a novel correlation clustering formulation of the quasispecies reconstruction problem which relies on semidefinite programming to accurately estimate the sub-species and their frequencies in a mixed population. Extensive comparisons with existing methods are presented on both synthetic and real data, demonstrating efficacy and superior performance of QSdpR.

## 1 Introduction

RNA polymerases that replicate viral genomes exhibit high error rates which cause relatively frequent point mutations in the viral genomic sequences. As a result, RNA viruses typically exist as collections of non-identical but closely related variants inside host cells. The diversity of viral populations, often referred to as viral quasispecies, adversely affects antiviral drug therapy and renders the vaccine design challenging [1], thus motivating their close studies. The quasispecies reconstruction (QSR) problem involves both the reconstruction of individual sequences in a population as well as the estimation of their abundances. Presence of sequencing errors in *next generation sequencing* (NGS) reads, short read lengths as well as small genetic distances between viral strains make QSR a hard problem to solve, even when sequencing coverage is high. Although conceptually similar to the single individual haplotyping problem, QSR has major additional challenges – the number of individual haplotypes is *a priori* unknown and the point mutations are in general poly-allelic rather than bi-allelic [2].

Recent approaches to solving the QSR problems include Bayesian inference methods such as ShoRAH [3] and QuRe[4], the non-parametric Bayesian approach based on Dirichlet process mixture model in [5] named PredictHaplo, Hidden Markov model based Quasirecomb [6], max-clique enumeration technique on read alignment graphs called HaploClique [2], graph-coloring based heuristic named VGA [7], and the reference assisted *de-novo* assembly reconstruction method named ViQuaS [8]. Generally, these methods can be categorized as read-graph based [2], [7], probabilistic inference based [3]–[6] and *de-novo* assembly based techniques [8]. The read-graph based methods [2], [7] rely on a combinatorial approach to analyze a graph with vertices that represent reads and edges that connect nodes corresponding to reads which overlap. Specifically, [2] formulates the QSR problem as that of enumerating the maximal cliques in the aforementioned graph, while [7], [10] map it to the coloring of graph vertices using structural constraints. In probabilistic methods, one formulates QSR as the problem of inferring hidden variables that model abundance of viral variants or point mutations and recombinations. In particular, PredictHaplo [5] uses an infinite mixture model to determine the number of species and reconstructs each haplotype by maximizing likelihood of the observed reads. Lastly, [8] performs local haplotype reconstruction as a reference-assisted *de-novo* assembly and arrives at a global solution by seeking overlap agreement of the locally reconstructed haplotypes. Moreover, quasispecies reconstruction methods often employ high-fidelity sequencing protocols [7] and barcode-tagging of genomes [9].

Building upon the method in [11] which approximately solves a semi-definite programming relaxation of the max-*K*-cut problem to perform single individual haplotyping, in this paper we extend the correlation clustering framework to viral quasispecies reconstruction. Specifically, we develop an accurate and computationally efficient algorithmic scheme that detects the number of strains in a population, reconstructs their genomes, and determines their frequencies. To ensure computational efficiency, the scheme removes spurious read overlaps with a negligible loss of accuracy. We test the performance of the proposed method on data sets emulating varying frequency spectrum, coverage and nucleotide diversities as well as on a widely used real data set introduced in [12] that contains Illumina MiSeq NGS reads from a mixture of 5 known HIV-1 strains present at non-uniform proportions. Benchmarking of the performance has been conducted in terms of the minimum error correction (MEC) scores [13], Reconstruction Proportions, Reconstruction Errors and Frequency Deviation errors. The results demonstrate superior performance of the proposed method as compared to state-of-the-art techniques including PredictHaplo, ShoRAH and ViQuaS. An open source software implementation of the method is freely available at https://sourceforge.net/projects/qsdpr.

The paper is organized as follows. In Section 2, we present an end-to-end correlation clustering framework for viral quasispecies reconstruction that includes detection of the number of species in a mixture. In Section 3, metrics used to assess performance of the proposed method are defined. Section 4 overviews both the synthetic and experimental data used for benchmarking, while Section 5 presents results of the benchmarking tests and the comparison with competing methods. Section 6 summarizes and concludes the paper.

## 2 Method

### 2.1 System Model

Let 𝒬 = {*q_k_*, *k* = 1,…,*K*} denote the set of *K* quasispecies strains present in a heterogeneous mixture of viruses. The strains in 𝒬 are strings of identical lengths consisting of alleles (i.e., nucleotides *A, C, G* and *T*), and differ from each other at a number of variant sites (i.e., at the locations of *single nucleotide polymorphisms* or SNPs). For simplicity, we restrict our attention to the mutations, i.e., nucleotide substitutions, and do not explicitly model insertions/deletions. Let *R* = {*r_i_*,*i* = 1,…, |R|} denote the set of overlapping reads aligned to a given reference genome; the reads are DNA fragments obtained via the so-called *shotgun* sequencing strategy used by next-generation sequencing platforms and effectively sample (with replacement) the strains in 𝒬. Sequencing platforms suffer from base calling errors which limit the reads to be much shorter than quasispecies strains. Note that the homozygous sites (i.e., sites containing alleles common to all strains) are not used in the quasispecies reconstruction and thus the corresponding bases can be removed from the reads covering those sites. Reads that cover one or no SNPs are also removed from the set since they are non-informative. Let there be 𝓁 variant sites that remain following the above pre-processing of sequencing data. Then, each *q_k_* can be thought of as a string of alleles of length 𝓁, while each read *r_i_* is a short, randomly positioned and potentially erroneous sub-string of one of the *q_k_*’s. The goal of viral quasispecies reconstruction is to segment this set of reads into as many clusters as there are viral strains (namely, *K*) so that each cluster consists of reads that sample one specific strain.

### 2.2 QSR as a Correlation Clustering Problem

Previously described clustering problem can be formalized by introducing a weighted read graph 𝒢 = (𝒱, 𝜀), where each vertex in 𝒱 corresponds to a read *r_i_* ∈ *R*, and each edge *e_ij_* ∈ *𝜀* represents the overlap between *r_i_* and *r_j_*. The weight or correlation associated with *e_ij_*, denoted as *ω_ij_*, is given by

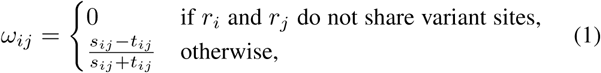

where *s_ij_* and *t_ij_* denote the number of matches and mismatches at the overlapping variant sites of *r_i_* and *r_j_*, respectively. Large *w_ij_* implies that *r_i_* and *r_j_* originate from the same strain while small *w_ij_* implies the opposite. Note that graph 𝒢 is highly sparse since, as pointed out in Section 2.1, the reads are much shorter than the genomic region of interest and thus each read overlaps with relatively few other reads. The edge weights defined in (1) have a direct implication on one of the properties of this partially observed graph, namely, signed weighted edge density is larger within clusters than across clusters, i.e., reads with higher correlation tend to be clustered together. The objective is then to cluster vertices *v* ∈ 𝒱 into *K* clusters such that the signed edge density within clusters is maximized, while the signed edge density across clusters is minimized. Note that sequencing errors in reads cause weights *ω_ij_* to deviate from true values, thereby making the graph clustering problem non-trivial. It has been shown in [16] that the non-sparse version of the correlation clustering problem is APX-hard.

In this paper, we approach viral quasispecies reconstruction by formulating it as a max-*K*-cut problem and solve its semidefinite relaxation akin to the approach to single individual haplotyping in [11]. The aim of max-*K*-cut is to partition a given set 𝒱 into *K* subsets 𝒱_1_, 𝒱_2_,…, 𝒱_*K*_ such that 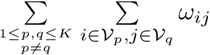 is maximized. A convex relaxation of the max-*K*-cut problem leads to a semi-definite program (SDP) which can, in principle, be solved using standard SDP solvers that rely on the interior point method. In particular, the resulting semidefinite problem is given by^1^

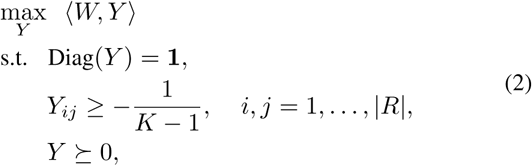

where *W* = {*ω_ij_*} is the edge weight matrix of G, 1 is the |*R*|-dimensional vector of all 1’s, and each row of the |*R*| × |*R*| matrix *Y* is a norm-1 vector which takes 1 of *K* possible values and has the property that its inner product with any other row gives −1/(K − 1) [17]. Such a definition of *Y* implies that its rank is *K*, which is then relaxed to rank |*R*| to arrive at the SDP formulation (2).

Viral quasispecies assembly with rare variants requires high data coverage and often leads to large-scale optimization problems. Since *Y* can be interpreted as a noisy version of an underlying matrix with data that originated from *K* clusters, it embeds a low dimensional rank-*K* structure within it. This is notable since SDPs with low-rank solutions can be solved efficiently [18]. Therefore, to find computationally feasible solutions to QSR, it is beneficial to express *Y* as *Y* = *VV^T^*, where *V* is a |*R*| × *K* matrix, and re-phrase optimization (2) in terms of this low dimensional matrix *V* (*K* ≪ |*R*|) such that 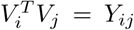. In particular, the Lagrangian relaxation of (2) is given by

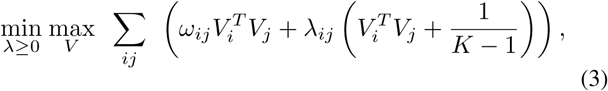

where λ = {λ_ij_} is an |*R*| × |*R*| matrix of Lagrange multipliers for the inequality constraints in (2). To solve (3), the objective function is alternately optimized with respect to *V*, keeping λ fixed, and then with respect to λ, keeping *V* fixed, via gradient descent. With λ fixed, the *i^th^* row of *V* is updated as *V_i_*←Σ_*j:e_ij_∈𝜀*_(*ω_ij_*+ λ_*ij*_)*V_j_*, followed by a normalization to preserve the unit norm property of rows; with *V* fixed, λ is updated according to 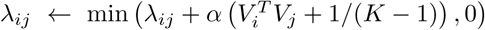 where *α* denotes the step size of the gradient descent algorithm and the factor multiplying α is the subgradient of the objective in (3) with respect to *λ*. Vectors of random entries are added as columns to *V* one at a time and the optimization of (3) is repeated until the rank of *V* becomes smaller than the number of its columns. The optimal solution *V_opt_* of this procedure will have a rank *r_opt_* ≥ *K*; therefore, to find a *K*-clustering of 𝒱 as in the original problem, a randomized projection heuristic is applied [17] - we project *V_opt_* onto an *r_opt_* × *K* matrix *P*, with *P_ij_* ~ 𝒩(0,1), and choose the *j^th^* cluster for *r_i_* if 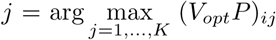. For each cluster, a consensus sequence of length 𝓁 is created by position-wise majority voting among those reads that fell within that cluster. This is followed by a greedy refinement step to further improve the overall objective function, that involves changing cluster memberships of reads as well as swapping alleles in the cluster consensus sequences. Finally, the consensus sequences are extended to full-length genomes by completing non-polymorphic (homozygous) sites with alleles from the reference genome to yield *K* quasispecies strains, whose frequencies are proportional to the size of corresponding clusters.

Deep sequencing of RNA virus strains, essential for detection of low frequency species in the mixture, results in a very high sequencing coverage (i.e., a very large read set *R*), leading to a high edge density in 𝒢. However, spurious read overlaps produce edges with uncertain weights which may induce errors in the quasispecies assembly solution. For example, if *r_i_* and *r_j_* are such that *t_ij_* = 0 (no mismatch) but *s_ij_* ≠ 0 (matches only), *ω_ij_* will be 1 irrespective of the actual value of *s_ij_*. Hence, as a graph sparsification step, we retain only those edges of 𝒢 in the clustering formulation which represent overlap of length above a certain threshold, i.e., edges that satisfy *s_ij_* + *t_ij_* ≥ *є*_o_, where *є*_o_ is an *edge overlap constant,* and set *ω_ij_* = 0 otherwise, thereby ensuring that the values of non-zero edge weights are relatively reliable. Moreover, edges characterized by comparable *s_ij_* and *t_ij_* (*ω_ij_* ≈ 0) are non-informative in terms of clustering; therefore, edges in 𝒢 are retained only if the edge weights satisfy |*ω_ij_*| ≥ *є*_*a*_, where *є*_*a*_ is an *edge ambiguity constant.* This leads to reduction in complexity and faster computation of the objective function in (3) without compromising quality of the clustering solution.

### 2.3 Determining the number of species

In order to reconstruct the quasispecies strains present within the viral population and infer their proportions, an assembly procedure needs to determine the number of clusters *K* into which vertices of 𝒢 are to be partitioned. A major challenge for most clustering methods is the requirement to pre-specify the number of clusters. Note that a clustering that relies on parsimonious cost objective functions (e.g., minimum error correction score) favors larger number of clusters over smaller (since the MEC score monotonically decreases in *K*). Therefore, such approaches may be conducive to overestimating the number of clusters, producing true clusters at the cost of a large number of false positives. The model selection problem in clustering is an open and active research topic and is known to be very difficult to solve. In this work, we quantify the quality of a clustering solution by using the Caliński-Harabasz criterion [19], also known as the *pseudo*-F index, defined as

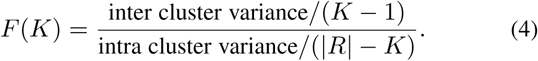

The terms in the numerator and denominator of (4) in the context of quasispecies reconstruction are defined as follows. Let *c*(*i*) ∈ {1,…, *K*} denote the index of the cluster containing *r_i_*, ∀ *i*. Let *n_k_* denote the number of reads in the *k^th^* cluster and *q_k_* denote the consensus of the reads in the *k^th^* cluster; moreover, let *q̅* be the consensus of *q_k_*, *k* = 1,…, *K*. If *H D*(·,·) denotes the Hamming distance between 2 strings over the alphabet {*A, C, G, T*}, counting non-gapped (i.e., sequenced) sites only, then the inter-cluster variance is defined as 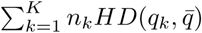 and the intra cluster variance is given by 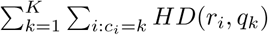. It is known that high values of the pseudo-F index indicate closely knit clusters [19], [20] and, in practice, the value of *K* for which this index is maximized is a good candidate for the number of underlying clusters. Therefore, the number of viral strains can be inferred by solving the quasispecies reconstruction problem over a preselected range of *K* and choosing the value of *K* for which the pseudo-F index has the highest value.

## 3 Performance metrics

We characterize the performance of viral quasispecies reconstruction algorithms by the accuracy of the number of reconstructed strains, the number of perfect (error-free) reconstructions among those strains, accuracy of the frequency estimation and the cumulative mismatch error of reads mapped to the reconstructed strains.

Let *n* and *N* denote the sizes of true and reconstructed quasispecies populations, respectively. Let *A* = {*a*_1_, *a*_2_,…, *a_n_*} and *B* = {*b*_1_, *b*_2_,…, *b_N_*} denote the two populations, and let 𝓁 be the length of the quasispecies strains. Assume *N* ≥ *n*. With *B*′ denoting the subset of *B* containing *n* strains with highest frequencies, define a function *f* : *A → B*′ that maps reconstructed strains to the true strains, i.e., for *i* = 1,…, *n*, *f* (*a_i_*) denotes the strain in *B*′ which is the closest neighbor to *a_i_* in terms of the Hamming distance. In order to ensure distinctive matching, *f* should be a one-to-one mapping. Hence, we choose *f* as

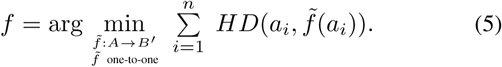

We further define a read mapping function *g*: *R* → *B′* as

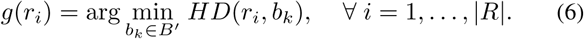

The cumulative mismatch error for the entire read set is referred to as the minimum error correction (MEC) score; the score expresses consistency between the observed reads and the reconstructed solutions [25]. It is formally defined as

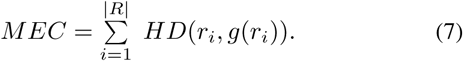

We define *Predicted Proportion* as the ratio of the reconstructed population size to the true population size, *N/n,* and *Reconstruction Proportion* as the ratio of the number of perfect (error-free) reconstructions to the total number of true species. A reconstruction is considered perfect or error-free if all alleles in the reconstructed strain match the corresponding alleles in one of the true species. To quantify accuracy of reconstruction, we define *Reconstruction Error* as

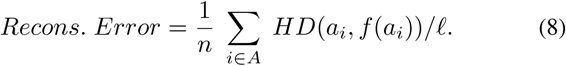

Lastly, accuracy of frequency estimation is measured by the total variation distance or 𝓁_1_-norm distance between the reconstructed and true quasispecies spectrum, i.e.,

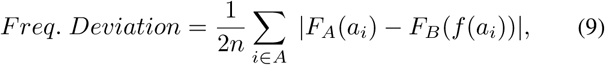

where *F_A_* : *A →* [0,1] and *F_B_* : *B′ →* [0,1] denote the frequencies of the true and reconstructed populations, respectively [26]. In the case when *N < n,* the metrics (8) and (9) are computed on a subset *A′* of *A* consisting of those strains which are the closest neighbors of the reconstructed strains, with *f* now defined as a mapping from *A′* to *B* in a manner similar to (5).

It is worthwhile reviewing the definition of nucleotide diversity, a measure of the genetic diversity present within a population. Specifically, for the set *A* given above, nucleotide diversity is defined as

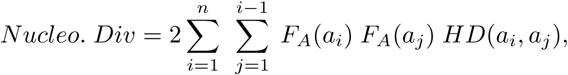

and similarly defined for *B*′.

## 4 Materials

### 4.1 Simulated data

In the first part of the simulation study, we synthesized datasets by emulating high-throughput next generation sequencing of viral populations, comprising of quasispecies strains present at uniform and non-uniform proportions and with varying number of species in the mixture. These datasets contain 2 × 350 bp (base pair) long paired-end^2^ reads at a coverage sufficiently high to facilitate reconstruction of the rarest species. First, we considered a mix of 5 viral strains and generated reads at an effective sequencing coverage 150*X*, where effective sequencing coverage is defined as the average number of sequenced bases per nucleotide position per quasispecies strain. Frequencies of the strains in the mix are assumed to be uniform, i.e., 20% for each strain. The strains are generated by introducing independent single nucleotide variations at uniformly random locations along the length of a randomly generated reference genome, at the rate of 1 mutation per 100 bases. To simulate realistic error rates of the Illumina’s NGS platforms, 1 base per 100 nucleotide positions is replaced with one of the remaining 3 bases. Inserts of the paired end reads are on average 1000 bp long with standard deviation of 100 bp. This dataset is referred to as *S*1. A second synthetic dataset, *S2,* contains reads that provide 200*X* coverage for 10 viral strains with uniform frequency distribution of the species (10% for each strain), where the mutation rate, sequencing error rate and insert length parameters are identical to those used in *S*1.

To emulate practical scenarios where viral populations are characterized by non-uniform proportions of the member strains, we generated two additional datasets of reads using identical parameters except for a varying frequency of strains. These two sets, denoted as *S*3 and *S*4, contain 5 and 10 strains and have effective coverage as 200*X* and 400*X*, respectively, sufficiently high to allow reconstruction of strains at frequencies as low as 5%.

The performance of QSdpR is also tested on a data set consisting of simulated reads generated by mimicking the characteristics of a real HIV-1 *in vitro* population, published in [12]. In particular, 5 quasispecies strains of length 1000 bp (typical length of a gene in the *pol* region of the HIV-1 genome) are simulated to have a SNP rate of 0.0986^3^. Paired-end reads of length 237 bp ±1% with 250 bp long inserts, typical of the dataset in [12], are simulated at an average coverage of 2000*X*; sequencing error rate is maintained identical to that in the previously discussed datasets. Frequencies of the quasispecies strains are set according to the values estimated using *Single Genome Amplification* on the *protease* gene [15] for the dataset in [12]. This data set is referred to as *S*5 and summarized in Table 1 along with all the other synthetic datasets.

**TABLE 1:** Summary of the description of simulated datasets

| Dataset | No. of Haplotypes | Nucleotide Diversity | Effective Coverage | Frequency (%) |
| --- | --- | --- | --- | --- |
| S1 | 5 | 0.3% | 150X | 20, 20, 20, 20, 20 |
| S2 | 10 | 0.5% | 200X | 10, 10, 10, 10, 10, 10, 10, 10, 10, 10 |
| S3 | 5 | 0.25% | 200X | 50, 25, 12.5, 6.25, 6.25 |
| S4 | 10 | 0.5% | 400X | 18, 16, 14, 12, 10, 8, 7, 5, 5, 5 |
| S5 | 5 | 3% | 2000X | 12.3, 9.2, 27.9, 38.4, 12.2 |

In the second part of the study, we synthesize datasets to assess the performance of QSdpR in settings where lengths of the quasispecies strains are varied from 1000 bp to 3000 bp. To this end, 2 × 150 bp long paired end reads with 200 bp inserts are simulated at an effective coverage of 500*X*; these reads sample a viral population of 5 quasispecies strains present at uniform proportions. The SNP rate and the sequencing error rate are identical to those in the simulation scenarios S1-S4. 10 datasets are simulated for each value of reference length and performance metrics averaged over the 10 runs are reported. These datasets are referred to as *L*1-*L*5.

### 4.2 Real data

To further benchmark the performance of QSdpR, we consider the HIV Five Virus Mix experimental dataset [12] and use the NGS reads generated by Illumina’s MiSeq sequencing platform. The data set consists of reads from a quasispecies population generated *in vitro* using 5 known HIV-1 strains named HIV-1 89.6, HXB2, JR-CSF, NL4-3 and YU2. The paired-end reads are on average 237 bp long and have standard deviation 26 bp; they are aligned to the HIV-1 *HXB2* reference genome and are obtained from Genbank (accession number SRP029432). Using state-of-the-art variant caller (see Section 4.3), 958 SNPs are observed along the whole length of the reference, out of which 690 SNPs are located within the various gene regions of interest. Sequencing depth for this data is highly non-uniform, as can be seen from Figure 1. Figure 1a shows the coverage in terms of sequenced bases per reference nucleotide position, while Figure 1b shows the coverage at the SNP locations only, along with the proportions of *A, C, G* and *T* bases at those locations. Figure 1b indicates that the quasispecies data is poly-allelic at several of the SNP sites. The ground truth for this quasispecies mixture is available at http://bmda.cs.unibas.ch/HivHaploTyper/, in the form of 5 individual strains that were sequenced independently before the mixture of viral strains was formed. Frequencies of the individual strains are estimated by amplifying *protease* gene of the *pol* region of the HIV-1 genome by means of *SGA* [15]. HIV-1 genome contains three major genes, namely, *gag* or the group specific antigen which codes for polyprotein, *pol* or polymerase which codes for viral enzymes such as reverse transcriptase (RT), integrase (int) and protease (PR), and *env* or “envelope”, which codes for glycoprotein, along with accessory regulatory genes such as *nef*, *vpr*, *vif* and *vpu*. Features of these gene regions are given in Table 2. The proposed quasispecies reconstruction method is applied to these genic regions for benchmarking of the algorithm performance [12][14].

**Fig. 1:**
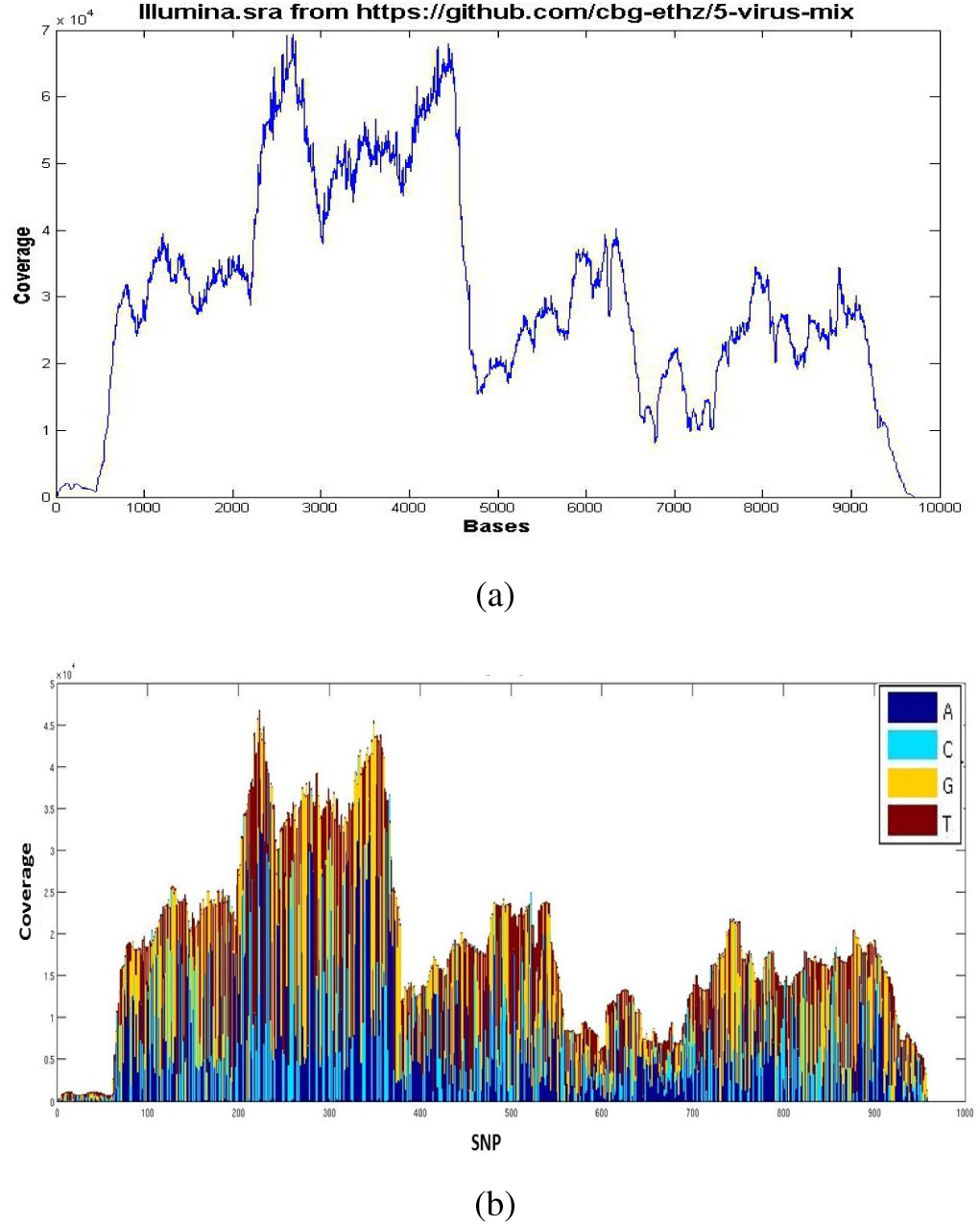
Coverage of the HIV Five Virus Mix dataset [12] in terms of the number of overlapping pair-end reads per base pair. This data, sequenced on Illumina MiSeq platform, is available at https://github.com/cbg-ethz/5-virus-mix. Subfigure (a) represents the raw read set before any quality clipping or filtering steps. Subfigure (b) represents the coverage and proportions of alleles at only the SNP locations, after pre-processing of read data. Lengths of the colored bars at each location denote proportions of A, C, G and T nucleotides as per read data.

**TABLE 2:** Lengths of the genomic regions (in base pairs), percentage nucleotide diversity and the number of single nucleotide polymorphisms (SNPs) for genes in the HIV-1 *Five Virus Mix* dataset. Nucleotide diversity is computed on the basis of the individual HIV-1 strains representing the ground truth.

| Gene | p17 | p24 | p2-p6 | PR | RT | RNase | int | vif | vpr | vpu | gp120 | gp41 | nef |
| --- | --- | --- | --- | --- | --- | --- | --- | --- | --- | --- | --- | --- | --- |
| Length (bp) | 396 | 693 | 413 | 297 | 1320 | 350 | 866 | 578 | 291 | 248 | 1533 | 1037 | 620 |
| Nucleo. Div.(%) | 2.08 | 1.04 | 1.15 | 1.04 | 1.05 | 1.81 | 0.94 | 1.88 | 1.71 | 3.15 | 2.71 | 2.27 | 2.66 |
| #SNPs | 46 | 45 | 32 | 19 | 87 | 37 | 49 | 61 | 31 | 31 | 32 | 133 | 87 |

### 4.3 Processing of the NGS datasets

The quasispecies reconstruction procedure starts with aligning the raw NGS reads to a given reference genome. Alignments were performed using ‘bwa mem’ algorithm (based on Burrows-Wheeler Transform) with the default settings [21]. Reads are filtered out if they are not mapped at all or not properly aligned to the reference, or if they are not primary alignments, PCR (*polymerase chain reaction*) duplicates or unable to pass quality filters (see [22] for aligned read flags). Reads having more than 2 consecutive ‘N’ basecalls are discarded. Next, the sites for multi-allelic variants in the aligned and sorted read set are detected by relying on a widely-used variant caller *FreeBayes*^4^ (version 0.9.20-16) [23]. Since *FreeBayes* requires ploidy information as input among other things, we set this parameter to a sufficiently high value (namely 25) in the experiments to ensure that the SNP diversity of the underlying strains is suitably captured. However, in the experiments, it is found that *FreeBayes* returns exactly identical set of SNPs for all values of ploidy ranging from 2 to 25. The reads are converted into the input fragment format [24], amenable to quasispecies reconstruction, using custom Python scripts. This format is an efficient representation of variant information suitable both for single-end and paired-end reads. It is to be noted that the software available at this time for conversion of aligned reads into fragment representation handles bi-allelic data only [24], which required us to write our own scripts to deal with multi-allelic variants. At the end of the core clustering procedure, results are converted into full-length quasispecies strains by inserting nucleotides from the reference genome into non-variant sites and the reads are re-aligned with the resultant strains. Frequency of a quasispecies strain is then evaluated based on the fraction of reads that are nearest to it in terms of sequence similarity or the Hamming distance. The software QSdpR takes aligned reads and variant information as inputs, along with a reference genome, and outputs set of quasispecies strains with corresponding frequencies.

## 5 Results and Discussions

Detailed results of the experiments on simulated and real data sets are presented in this section. For convenience, the quasispecies reconstruction technique described in this paper is labeled and referred to as QSdpR. Our software implementing the QSdpR algorithm is made available as open source software at the location https://sourceforge.net/projects/qsdpr.

### 5.1 Results on simulated data

We tested our proposed method on the synthetic data sets *S*1-*S*5 and characterized its performance in terms of the metrics introduced in Section 3 (we can evaluate these metrics since in simulations the true quasispecies strains are known). The mismatch errors (i.e., the MEC scores) are shown in Figure 2, while *Predicted Proportion, Reconstruction Error* and *Frequency Deviation* metrics are reported in Table 3. For a meaningful and fair MEC comparison in Figure 2, we have excluded the cases where the reconstructions are partial, i.e., the recovered sequences are not full lengths (because fewer reads would map to the partial sequences, leading to lower MEC score). We compared QSdpR performance with that of PredictHaplo [5], ViQuaS [8] and ShoRAH [3]. Among the considered schemes, our method achieves the lowest MEC scores for all of the data sets considered here, followed by PredictHaplo (PredictHaplo failed to run successfully on *S*2 even though the sequencing coverage is as high as 200*X*), while ViQuaS and ShoRAH have much higher mismatch errors in all cases. We attribute the superior MEC performance of QSdpR to the fact that the correlation clustering framework is explicitly concerned with minimizing the cumulative Hamming distance between the reads and the reconstructed strains. It is worthwhile pointing out that since in practice the ground truth for a quasispecies population is generally unavailable (discovering it is the entire purpose of QSR), performance metrics such as the *Reconstruction Error* and frequency mismatch cannot be computed in the experimental settings. In these scenarios, MEC score is a proxy measure of the quality of the reconstruction.

**Fig. 2:**
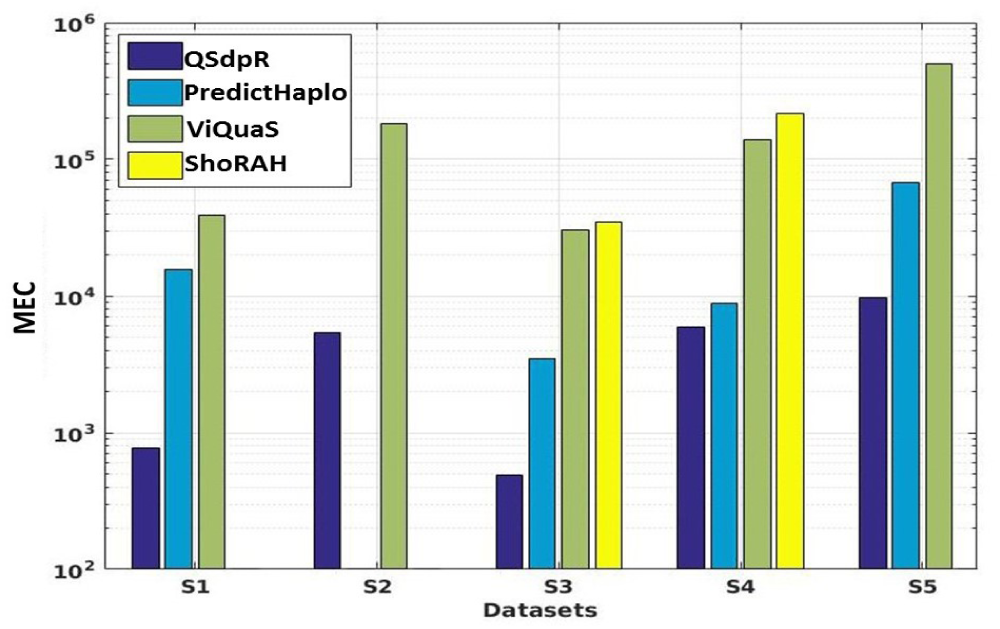
The MEC score comparison of QSdpR, PredictHaplo, ViQuaS and ShoRAH on the Simulated Datasets S1-S5. PredictHaplo did not run on S2. ShoRAH returned haplotypes with 72%, 44.6% and 93.6% of reference genome lengths on sets S1, S2 and S5, respectively.

**TABLE 3:** Performance evaluation of QSdpR on the simulated datasets *S*1-*S*5. QS, PH, VQ and SH denote QSdpR (correlation clustering), PredictHaplo, ViQuaS and ShoRAH, respectively. Boldface value in each row indicates the best performance for the given metric. PredictHaplo did not run on S2. ViQuaS reconstructed only one strain for S5, hence is excluded from the comparison.

| Dataset | Predicted Proportion | | | | Reconstruction Error ($\times 10^{-3}$) | | | | Frequency Deviation ($\times 10^{-2}$) | | | |
| --- | --- | --- | --- | --- | --- | --- | --- | --- | --- | --- | --- | --- |
|  | QS | PH | VQ | SH | QS | PH | VQ | SH | QS | PH | VQ | SH |
| S1 | **1** | **1** | 6.2 | 13 | **0** | 9.2 | 5.7 | 6.7 | **0.06** | 0.78 | 1.68 | 7.2 |
| S2 | **1** | NA | 8.5 | 13 | **0.4** | NA | 9.9 | 4.7 | **0.06** | NA | 1.28 | 3.62 |
| S3 | 1.2 | 0.8 | 6 | 9.6 | **4** | **4** | 6.5 | 6.6 | 0.09 | **0.03** | 8.2 | 5.21 |
| S4 | 1.09 | **1** | 4.3 | 16.7 | 7.3 | **4.7** | 10.2 | 5 | 1.33 | **0.88** | 2.78 | 3.43 |
| S5 | **1** | 0.8 | 0.2 | 56.4 | **0** | 0.25 | NA | 6.6 | **0.01** | 3.07 | NA | 8.96 |

From Table 3, it can be seen that QSdpR infers the number of underlying strains correctly for 3 out of the 5 data sets, namely for *S*1, *S*2 and *S*5, as indicated by the Predicted Proportion values. For datasets *S*3 and *S*4, it overestimates the number of quasispecies strains by 1. On the other hand, PredictHaplo underestimates the number of species by 1 for sets *S*3 and *S*5, and infers it correctly for *S*1 and *S*4. ViQuaS and ShoRAH significantly overestimate this quantity (except for the set *S*5 where ViQuaS reconstructs only one mixture component). In terms of *Reconstruction Error*, our method is able to recover each of the true strains without a single base mismatch for *S*1 and *S*5; for *S*3, it matches the performance of PredictHaplo, which does not provide error-free reconstruction in any of the data sets on which it could successfully run. However, in set *S*4, PredictHaplo provides the smallest recovery error (precisely, it makes 26 nucleotide mismatches fewer than our method). ShoRAH has better reconstruction than our method only on *S*4. While our correlation clustering technique provides the most accurate spectra reconstruction for *S*1 and *S*5, it is less accurate than PredictHaplo on sets *S*3 and *S*4. This is due to overestimation of the number of strains for these 2 sets, which leads to misclassification of some reads (i.e., they are assigned to erroneously created clusters) thus causing discrepancy between the inferred frequencies and the correct ones. The latter effect is much more pronounced in ViQuaS and ShoRAH, primarily because they significantly overestimate the number of species.

To validate the efficacy of the approach to inferring the number of quasispecies strains as outlined in Section 2.3, we show normalized *F*(*K*) in Figure 3 for the synthetic data sets *S*1, *S*3 and *S*5 and all the 13 genes of the experimental dataset. Note that the true number of quasispecies strains in each of the considered datasets is 5. It is evident from the figure that the correct value of *K* maximizes the normalized pseudo F statistics for *S*1 and *S*5 (except for *S*3, where the metric is maximized by choosing *K* = 6, leading to an overestimation of the number of species in the mixtures), and for all the HIV-1 genes (except *PR* and *nef* genes, where the metric is maximized by choosing *K* = 7 and *K* = 4 respectively).

**Fig. 3:**
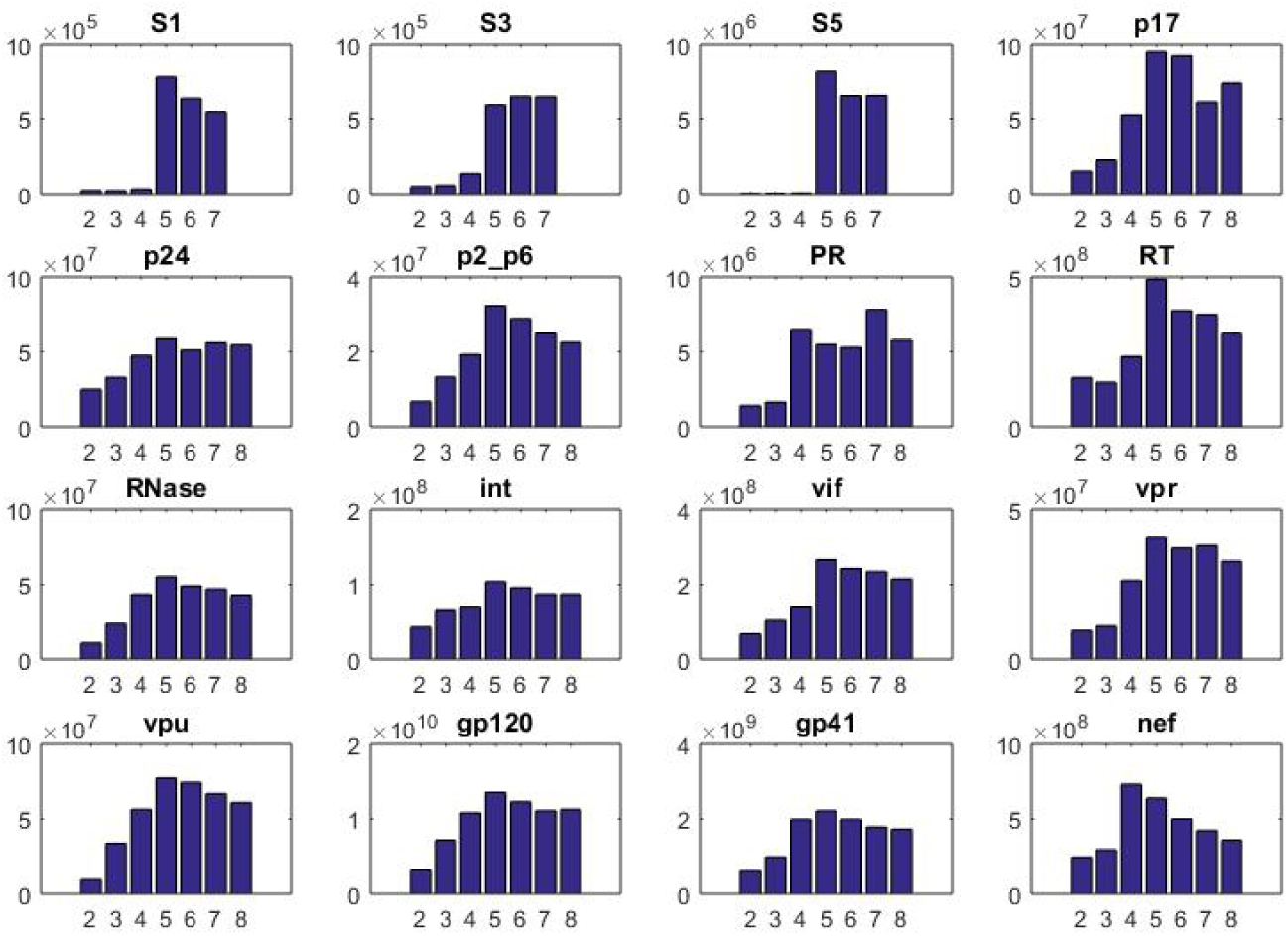
Normalized Pseudo F statistics as a function of the parameter *K* for simulated data sets *S*1, *S*3, *S*5 and 13 HIV-1 genes *p*17 through *nef*. The true number of species for each dataset is 5. Value of *K* is correctly inferred for *S*1 and *S*5 among simulated sets and for all HIV-1 genes except *PR* and *nef*.

To demonstrate how the running time of QSdpR scales with parameter *K*, in Figure 4 we show runtimes for the values of *K* ranging from 2 to 10 for simulation datasets *L*1 to *L*4. QSdpR is applied to each of these sets and the resulting running times averaged over 10 simulation runs for each set are plotted in Figure 4. Since pre-processing of reads is common for all *K* and post-processing (i.e, the construction of full length quasispecies strains from clustering solutions) time is consistent across *K*, Figure 4 shows only the runtimes of the procedure outlined in Section 2.2. This figure indicates that the computational overhead needed to determine the number of species according to the procedure outlined in Section 2.3 is practically feasible.

**Fig. 4:**
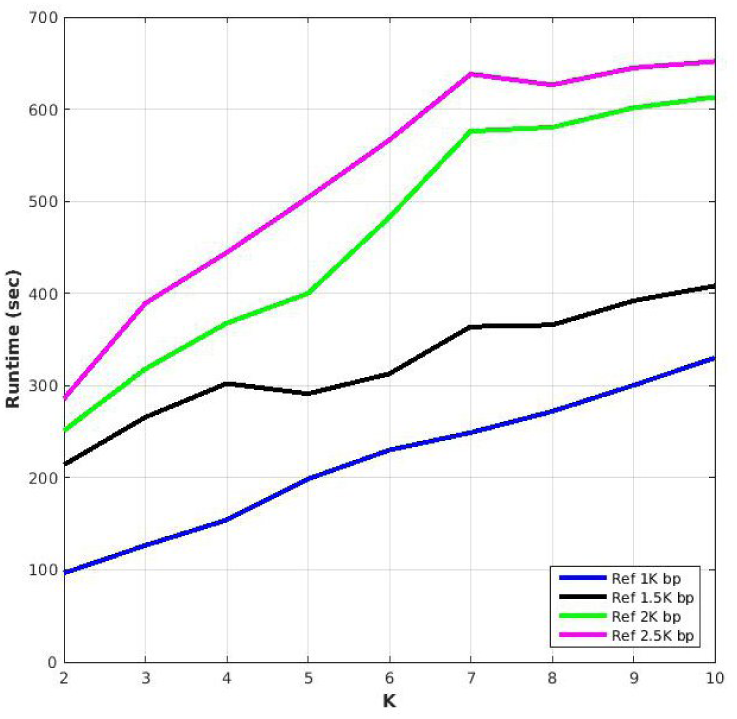
Comparison of QSdpR runtime as a function of the parameter *K* for simulation sets *L*1-*L*4. For the sake of simplicity, time required for post-processing of reads is not included in runtime calculation. The values shown are based on average of 10 Monte Carlo runs of simulation sets for each *K* and reference lengths.

In Figure 5, the runtimes of the proposed method are compared with those of the competing methods including PredictHaplo and ViQuaS using the simulation sets *L*1-*L*4. The runtimes shown in the figure are averaged over the 10 runs for each of the methods. It can be seen that the runtime of QSdpR is fairly comparable to that of PredictHaplo; in particular, the runtimes of the two algorithms are very close to each other for the case of 1000 and 1500 bp long quasispecies strains. The difference in runtime, however, is more prominent for longer strains. On the other hand, QSdpR is much faster than ViQuaS for all of the considered lengths of quasispecies strains. Note that we did not include the performance of ShoRAH here since it is the slowest of the four methods and its runtime distorts the scale in Figure 5. It is worthwhile pointing out that the slightly faster performance of PredictHaplo comes at a cost of underestimating the number of quasispecies strains present in the mixture. Indeed, the number of strains inferred by PredictHaplo for the above setup (average of 10 simulation runs) is given by 1, 2, 1.5 and 2, respectively, for sets *L*1 through *L*4, while the numbers of strains inferred by the proposed method is 6, 7, 8 and 10. The corresponding numbers returned by ViQuaS are 57, 100, 137 and 163.5, respectively. Clearly, PredictHaplo is unable to infer the richness of the diversity of viral populations in all the cases, with the minimum deviation from true number of species (5) being 3. QSdpR, on the other hand, infers the most accurate *K* for strain lengths 1000 bp (6 species) and the deviation increases with longer strain lengths. ViQuaS suffers from a gross overestimation of the number of species in all of the considered cases.

**Fig. 5:**
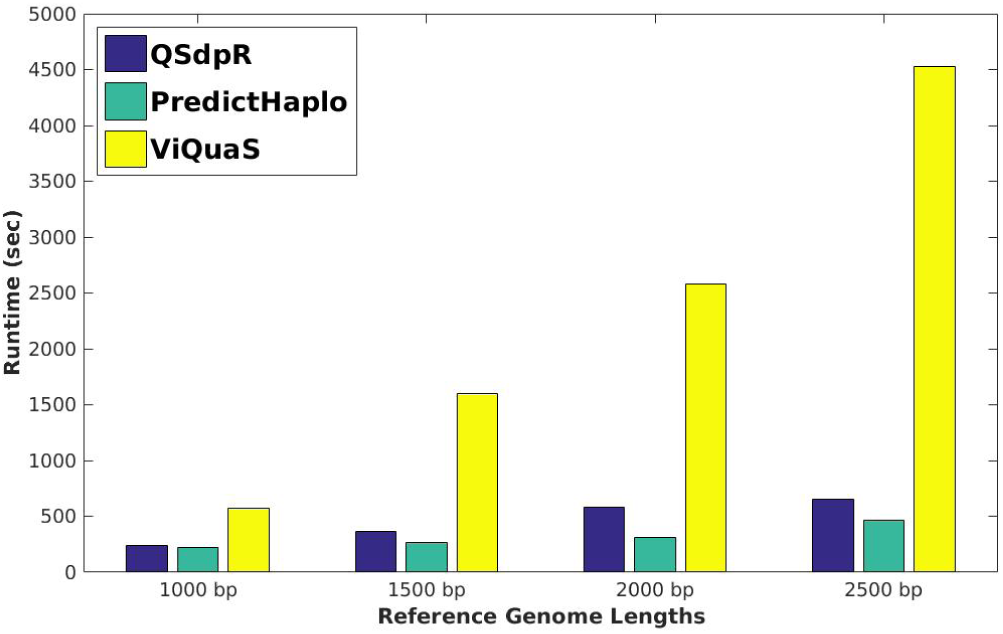
A comparison of runtime of QSdpR with PredictHaplo and ViQuaS for the simulation sets *L*1 to *L*4. The values shown are based on the average of 10 Monte Carlo runs of simulation sets for each method. ShoRAH is not included in the comparison since it is slowest among all the methods considered.

In Figure 6, we study how the performance of the proposed method depend upon the values of the parameters *є*_o_ and *є*_*a*_. Recall that a higher value of either of these constants leads to an increased removal of spurious (and unreliable) edges from the correlation graph, resulting in a decrease in runtime complexity. In Figure 6, reconstruction error percentage rate and runtime of QSdpR are plotted against a range of values of *є*_o_ and *є_a_* for the simulation sets *L*3 and *L*5, corresponding to strains of length 2000 and 3000 base pairs respectively. The metrics for each of the simulation scenarios have been averaged over 10 sets. For the sake of simplicity of the benchmarking experiment, we assume prior knowledge of *K*. As can be seen from the subfigures of Figures 6, the running time decreases as the read graph becomes sparser with increasing *є*_o_ and *є_a_*. At the same time, the reconstruction accuracy improves since the clusters of reads corresponding to each quasispecies now contain a progressively higher fraction of reliable overlaps between the reads.

**Fig. 6:**
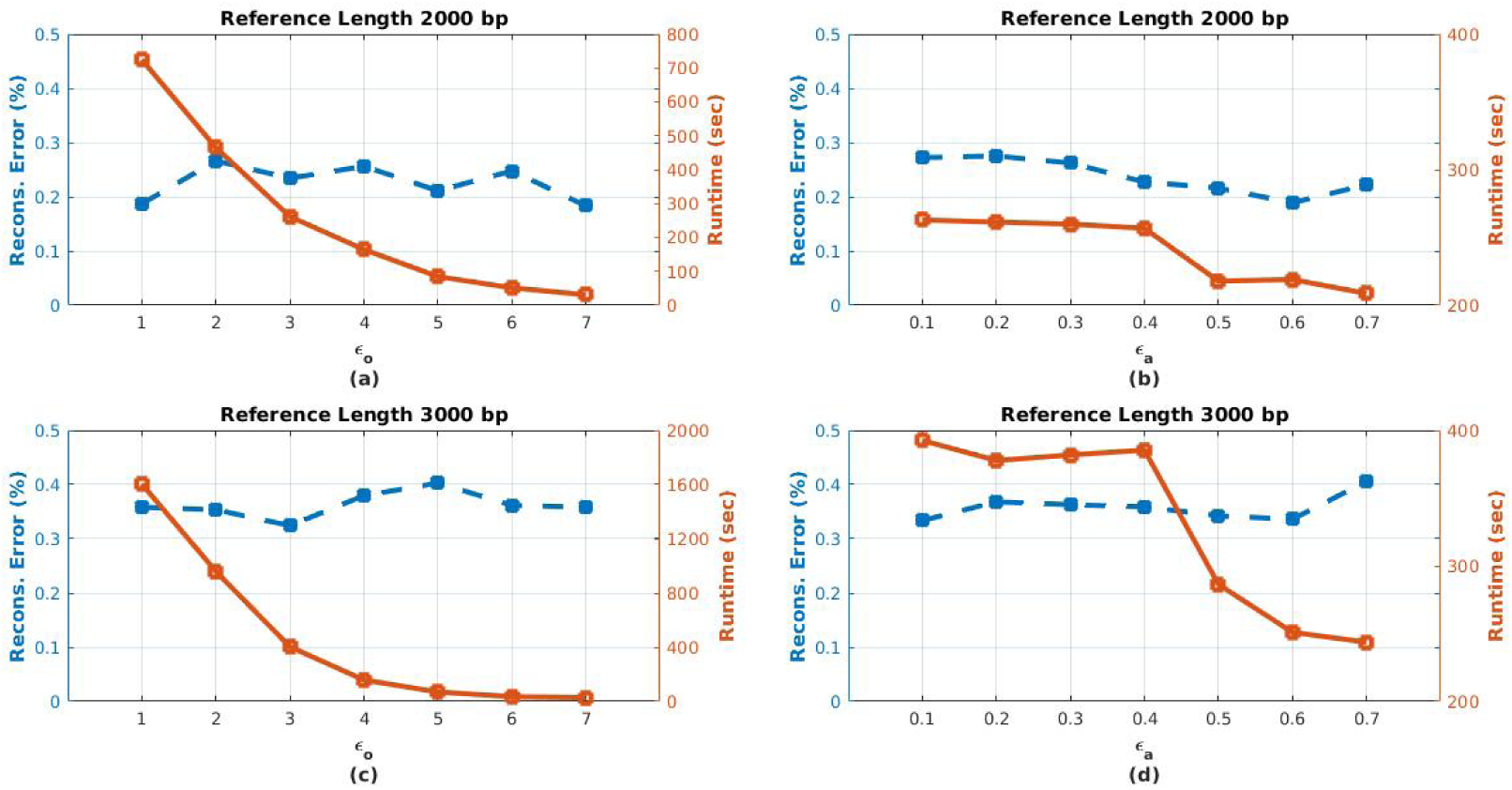
Performance variation of QSdpR with the correlation graph sparsification parameters, namely, edge overlap constant є_o_ (Figures 6(a),(c)) and edge ambiguity constant є_a_ (Figures 6(b),(d)) for quasispecies strains of lengths 2000 base-pairs (set *L*3, Figures 6(a),(b)) and 3000 base-pairs (set *L*5, Figures 6(c),(d)) The solid lines in each plot denote the running time of the proposed algorithm and the dashed lines in each plot denote the percentage reconstruction error. The values shown have been averaged over 10 simulation sets for each value of є_o_ and є_a_. є_a_ is fixed at 0:1 for Figures 6(a),(c) and є_o_ is fixed at 3 for Figures 6(b),(d).

Note that, however, as the graph grows sparser the clusters grow smaller in size and thus the effective coverage drops; this may lead to unreliable reconstruction of consensus strains at the end of the clustering procedure. As a result, the reconstruction error rate of the overall method may degrade. Moreover, higher values of *є*_o_ and *є*_a_ do not necessarily lead to a monotonic decrease in the complexity of the algorithm. A good practice is to choose those value of the constants which minimize the reconstruction error rate over a wide range of settings. For example, in Figure 6(c), *є*_o_ = 3 is the most favorable choice while in Figure 6(b), having *є_a_* = 0.5 optimizes the error rate over the given range.

### 5.2 Results on the experimental data

In this section, we report the results of comparing performance of the proposed method with competing algorithms when applied to reconstructing HIV-1 *Five Virus Mix* data set. Gene-wise quasispecies reconstruction is performed on the major genic regions of this single strand RNA genome; the performance metrics are computed for each of these regions. In order to determine the value of *K* used in reconstruction, we analyzed the 4036 bp long genomic region of the HIV-1 genome encompassing the *gag-pol* region. For cross-verification, we repeated the procedure for finding *K* with all the 13 individual gene regions and in 11 cases obtained the correct number of clusters (see Figure 3). Performance of the proposed method applied to gene-wise reconstruction is compared to that of PredictHaplo and ShoRAH. ViQuaS could not be used for gene-wise reconstruction since the current version of that software does not support specifying genomic regions while upon trying to run it for genome-wide reconstruction, the program did not complete its run in 36 hours on an 8-core machine. Two other recent approaches, namely *Haploclique* [2] and *VGA* [7] could not be used due to code execution problems and lack of active development support from the authors.

Figure 7 shows the MEC score comparison of our correlation clustering method with that of PredictHaplo and ShoRAH. QSdpR achieves better MEC scores than both PredictHaplo and ShoRAH for all 13 genes of HIV-1^5^. The performance is further characterized by *Predicted Proportion, Reconstruction Proportion, Reconstruction Error* and *Frequency Deviation,* all summarized in Table 4. As can be seen from this table, *Predicted Proportion* of our method is best among all 3 methods for all 13 genes, while that of ShoRAH is the worst. *Reconstruction Proportion* of QSdpR is better than that of PredictHaplo for 5 genes and equal to that of PredictHaplo for 3 genes. Compared to ShoRAH, QSdpR is better for 9 out of 13 genes. In particular, QSdpR is able to recover all 5 true quasispecies strains in 3 out of 4 genes in the *pol* region comprising of *PR, RT*, *RNase* and *int* genes. It is interesting to note that QSdpR in general has higher *Reconstruction Proportion* for genes with lower nucleotide diversities (see Table 2). In terms of *Reconstruction Error,* QSdpR outperforms PredictHaplo in 6 of the genes while for the remaining ones PredictHaplo has a better performance. As for ShoRAH, QSdpR has better performance on 9 of the genes and comparable performance in 1 gene. Since the proposed method relies only on SNPs to distinguish viral strains in a population, its *Reconstruction Error* may be higher in the genomic regions which harbor a significant portion of non-substitution mutations in the HIV-1 genome. Finally, in terms of *Frequency Deviation*, our method has equal or better performance than PredictHaplo on 8 genes and better than ShoRAH on 11 genes. The genes where other methods achieve slightly better performance are those in the gapped portion of the genome.

**Fig. 7:**
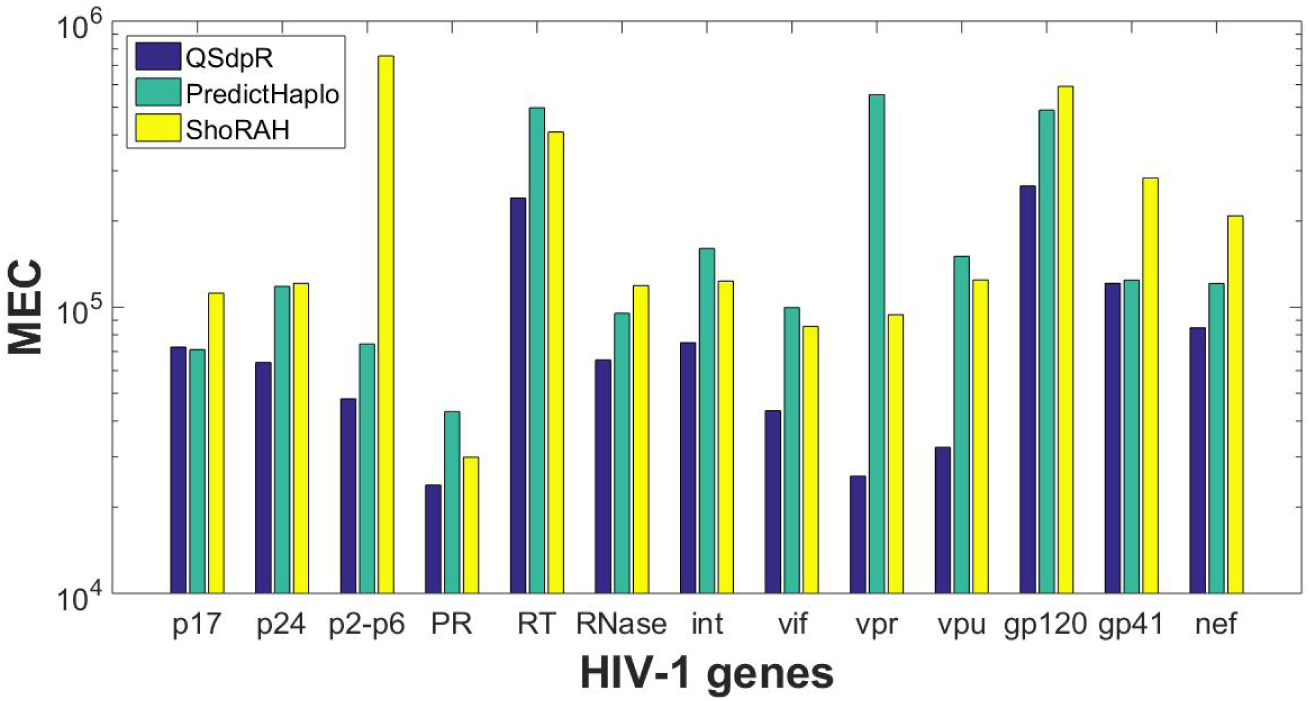
The MEC score comparison of QSdpR, PredictHaplo and ShoRAH on the HIV-1 *Five Virus Mix* dataset.

**TABLE 4:**
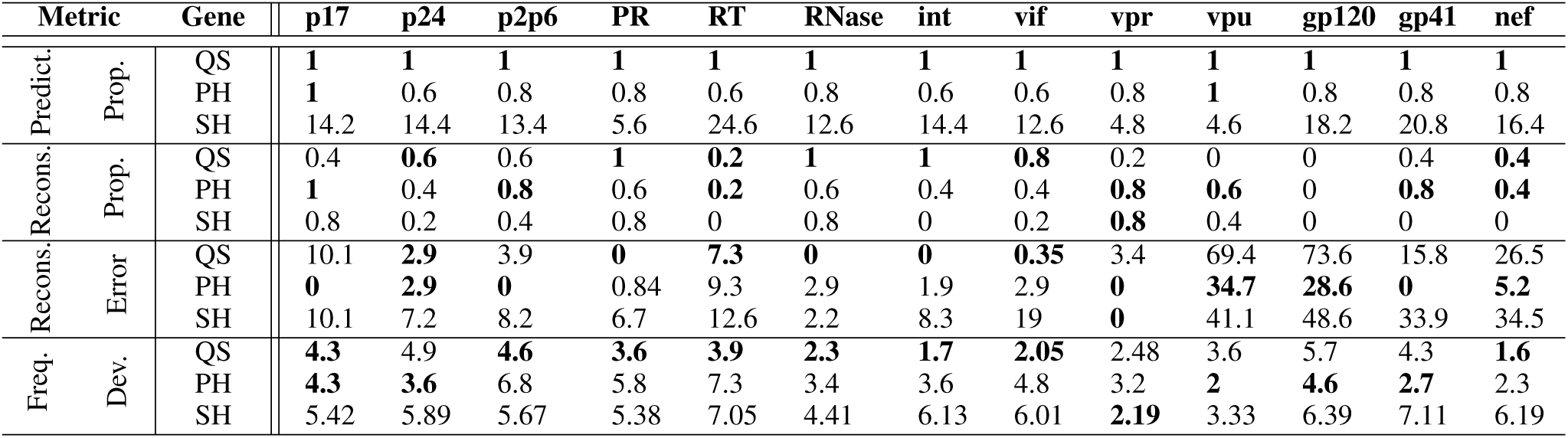
Comparison of *Predicted Proportion, Reconstruction Proportion, Reconstruction Error* and *Frequency Deviation* on the real HIV-1 *Five Virus Mix* data. QS, PH and SH refer to QSdpR (correlation clustering), PredictHaplo and ShoRAH, respectively. *Reconstruction Error* is to be multiplied with 10^−3^ and *Frequency Deviation* error is to be multiplied with 10^−2^ to get the actual numeric value. Boldface value in each column indicates the best performance for the given metric in that column.

## 6 Conclusion

Inference of RNA viruses in heterogeneous mixtures and estimation of their relative proportion within the quasispecies has been an active area of research in recent years. In this paper, we proposed QSdpR, an end-to-end framework for viral quasispecies reconstruction based on a correlation clustering formulation of the problem. The clustering is relaxed to a convex optimization problem and efficiently solved exploiting underlying sparse structure of the solution. We tested the method on synthetic data with uniform and non-uniform quasispecies spectra and varying diversity and mutation rate conditions. Moreover, the method was also tested on a widely known experimental HIV-1 dataset having 5 known strains. In all the scenarios, the proposed method competes favorably with the existing methods, providing accurate estimation of the viral quasispecies spectrum.

## Footnotes

1 〈*A*,*B*〉 denotes matrix dot product, given by 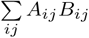 for matrices *A* and *B*.

2 Paired-end reads consist of two segments (a *pair*) that are separated by “inserts” or regions of unknown content but known length statistics.

3 SNP rate is inferred by analyzing haplotypes in the ground truth data.

4 However, QSdpR is not restricted to any particular choice of variant callers

5 except p17 gene, where MEC scores of QSdpR is comparable to Predic-tHaplo

